# Loss of Shp1 impairs myeloid cell function and causes lethal inflammation in zebrafish larvae

**DOI:** 10.1101/2022.06.23.497321

**Authors:** Maaike Allers, Petra A. Bakker, Herman P. Spaink, Jeroen den Hertog

## Abstract

*PTPN6* encodes SHP1, a protein tyrosine phosphatase that has an essential role in immune cell function. SHP1 missense and splice site mutations are associated with neutrophilic dermatoses and emphysema in humans, which resembles the phenotype seen in mice that lack functional SHP1 partially. Complete lack of SHP1 function in mouse *motheaten* mutants leads to severe defects and lethality at 2-6 weeks after birth. Here, we investigated the function of Shp1 in developing zebrafish embryos. We generated a *ptpn6* knockout zebrafish line lacking functional Shp1. Zebrafish develop ex utero and are transparent, which facilitates analysis of the function of Shp1 during early hematopoietic development. Shp1 knockout caused severe inflammation and lethality around 17 days post fertilization (dpf). During early development the myeloid lineage was affected, which resulted in a decrease in the number of neutrophils, and a concomitant increase in the number of macrophages. The number of emerging hematopoietic stem and progenitor cells (HSPCs) was decreased, but due to hyperproliferation, the number of HSPCs was higher in *ptpn6* mutants than in siblings at 5 dpf. Finally, directional migration of neutrophils and macrophages was decreased in response to wounding and less macrophages were recruited to the wound site. Yet, regeneration of the caudal fin fold was normal. We conclude that loss of Shp1 impairs neutrophil and macrophage function and caused severe inflammation and lethality at the larval stage.

## INTRODUCTION

The non-receptor protein tyrosine phosphatase SHP1 encoded by the *PTPN6* gene is a key regulator of immune cell function. SHP1 consists of two N-terminal Src-homology 2 (SH2) domains, a catalytic domain and a C-terminal regulatory tail, and is mainly expressed in hematopoietic cells (Neel, Gu and Pao, 2003). Because of its important role in regulating immune cell function, SHP1 has become an interesting target for treatment of auto-immune diseases and cancer in recent years (Watson *et al*., 2016).

SHP1 function has been studied extensively in the context of the *motheaten* (*me/me*) mouse that has a spontaneous recessive mutation in *Ptpn6*. This mutation creates a cryptic splice site, which results in the loss of functional SHP1 (Tsui *et al*., 1993). The homozygous mutation leads to immune deficiency, widespread inflammation, skin lesions and death within 2- 6 weeks due to lethal pneumonitis characterized by infiltration of myeloid cells (Green and Shultz, 1975). In the decades following the initial identification of the *me/me* mouse, multiple alternative less lethal mutations in *Ptpn6* have been described. *Motheaten viable* has a mutation in a splice consensus site of *Ptpn6*, which results in alternative splicing. SHP1 from *motheaten viable* mice exhibits 80% reduction in phosphatase activity and *motheaten viable* mice are lethal at 9-12 weeks (Shultz *et al*., 1984). *Spin* has a Y208N mutation in the C-terminal SH2 domain of SHP1, resulting in 50% residual catalytic activity and *spin* mice die after more than one year (Croker *et al*., 2008). The symptoms of *motheaten viable* are highly similar to the symptoms of *motheaten*, although they develop lethal pneumonitis approximately 8 weeks later. *Spin* mutants do not show lethal pneumonitis or immunodeficiency. This is likely due to the higher residual phosphatase activity of mutant SHP1 in *spin* mice. Heterozygous missense and splice variant mutations in *PTPN6* have also been found in human patients. These mutations are associated with emphysema and neutrophilic dermatoses (Nesterovitch *et al*., 2011; Bossé *et al*., 2019).

Compound mouse knock out lines and conditional knock out strains were generated to study the function of depletion of SHP1 in specific cell types. Mouse double-knockouts lacking RAG1 and SHP1 still show the *motheaten* phenotype, which indicates that the acquired immune system is not essential for the symptoms caused by loss of SHP1 (Yu *et al*., 1996). Conditional neutrophil-specific knockout of SHP1 does not recapitulate the lethal pneumonitis phenotype nor autoimmunity that was observed in the complete SHP1 knockout. Yet, knockout of SHP1 in the neutrophil specific lineage causes dermal inflammation. Knockout of SHP1 in the macrophage specific lineage does not cause inflammation, but it does cause lymphadenopathy and autoimmunity (Abram *et al*., 2013; Abram and Lowell, 2017), indicating that depletion of SHP1 in different cell types results in distinct hematologic defects.

Zebrafish do not have a functional acquired immune system until the larvae are 4-6 weeks old (Lam *et al*., 2004). During early development, zebrafish only have innate immunity. In addition, zebrafish provide the opportunity to investigate embryonic development from the start, due to their translucent eggs and embryos. Therefore, zebrafish is the ideal model system to investigate the function of Shp1 in the innate immune system in a whole organism.

Previous morpholino-mediated knockdown studies in zebrafish embryos showed that Shp1 knockdown reduced the ability of embryos to combat infection with *Salmonella typhimurium* and *Mycobacterium marinum*. The innate immune system was hyperactivated to a contra-productive level. Experiments suggest that Shp1 functions as a negative regulator that imposes a tight control over the level of innate immune response activation (Kanwal *et al*., 2013).

Here, we developed a genetic zebrafish model lacking functional Shp1 to study the hematopoietic system during early development in the absence of Shp1. Zebrafish mutant for Shp1 recapitulated the lethality and inflammation seen in *motheaten* mice. In addition, we investigated the development of HSPCs and all major blood lineages. We found that macrophage numbers were increased, whereas neutrophil numbers were decreased during early development. Emergence of HSPCs was reduced, but their subsequent proliferation was strongly increased. Finally we show that recruitment of macrophages and neutrophils to wound sites is disturbed in Shp1 mutants. Our results indicate that Shp1 has a role in hematopoietic development from the start until the development of specific cell types of the myeloid lineage, and Shp1 has an essential role in myeloid behavior after development as well.

## RESULTS

### Shp1 knock-out leads to inflammation and is lethal at late larval stage

We generated mutations in the zebrafish *ptpn6* gene using CRISPR-Cas9 technology at the 1 cell stage, followed by screening of the F0 generation for germline mutations. We identified a 7bp deletion in exon 4 resulting in a frameshift and a premature stop codon 10 amino acids downstream of the mutation site. Exon 4 encodes the C-terminal part of the N-terminal SH2 domain positioning the mutation well up-stream of the catalytic PTP domain and predicts production of a severely truncated protein of 96 amino acids (Fig 1A). To confirm the absence of Shp1 protein we detected total Shp1 protein levels in 5dpf zebrafish embryo lysates with a zebrafish Shp1 specific antibody, which we generated (Fig 1B). At 5dpf *ptpn6* mutants show no obvious phenotype (Fig. S1), in contrast to previous morpholino studies, which showed pleiotropic defects from 2 dpf onwards (Kanwal *et al*., 2013).

**Figure 1.**
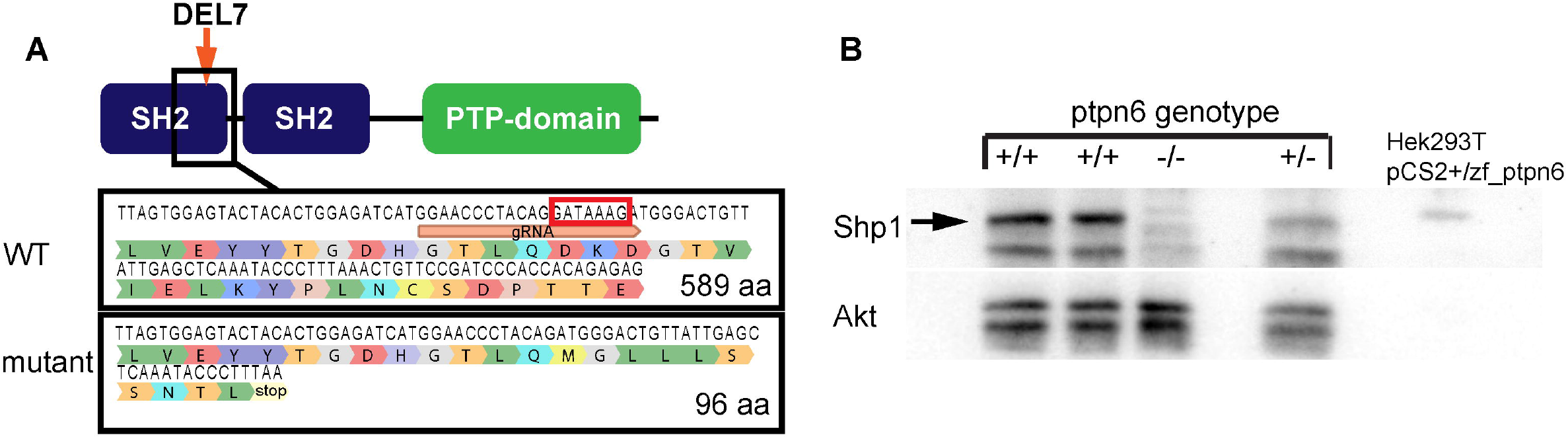
Zebrafish *ptpn6* mutant generated using CRISPR/Cas9 is a strong null allele. (A) Schematic overview of Shp1. gRNA was targeted to the C-terminal end of the N-terminal SH2 domain, inducing a 7 bp deletion in exon 4, indicated by the red box in the wild type sequence. Mutation was confirmed by sanger sequencing after sub-cloning of F1 DNA. The 7 bp deletion leads to a frameshift and a stop codon in exon 4, 10 amino acids downstream of the mutation site. (B) Immunoblot detecting endogenous Shp1 in WT and heterozygous embryos, but not homozygous *ptpn6* mutant embryos. Lysate of transfected Hek293T cells expressing zebrafish Shp1 was used as a control for detection of zebrafish Shp1. Akt-specific antibody was used to monitor equal loading.

During later larval stages the *ptpn6* mutants appear smaller and skinnier than siblings, and develop a curved phenotype. Moreover, as described for the *motheaten* mouse, the mutants show abnormalities in their skin epithelium (ruffled epithelial edges and bumps) (Fig 2A-D) (Green and Shultz, 1975). To look at infiltration of the skin with immune cells we used the *Tg(mpx:eGFP)* line marking myeloid-specific peroxidase producing neutrophils. We followed the developing larvae, mutants and siblings, over time and imaged them either at the moment of sacrifice or at the end point of the experiment (either 12 or 19dpf). Surprisingly, we did not only observe neutrophil infiltration of the skin, but the most prominent accumulation of neutrophils was found in the gill area (Fig 2C,D). *Motheaten* mice succumb to pneumonitis which is characterized by neutrophil and macrophage accumulation in the lungs. We semi-quantified the neutrophil accumulation phenotype by scoring the number of neutrophils in the gill and mandibular area in three categories: normal (WT), elevated, and high, at 7-12 dpf or 13-19 dpf (Fig 2E,F). It is evident that neutrophil numbers in the scored area are greatly increased in mutants.

**Figure 2.**
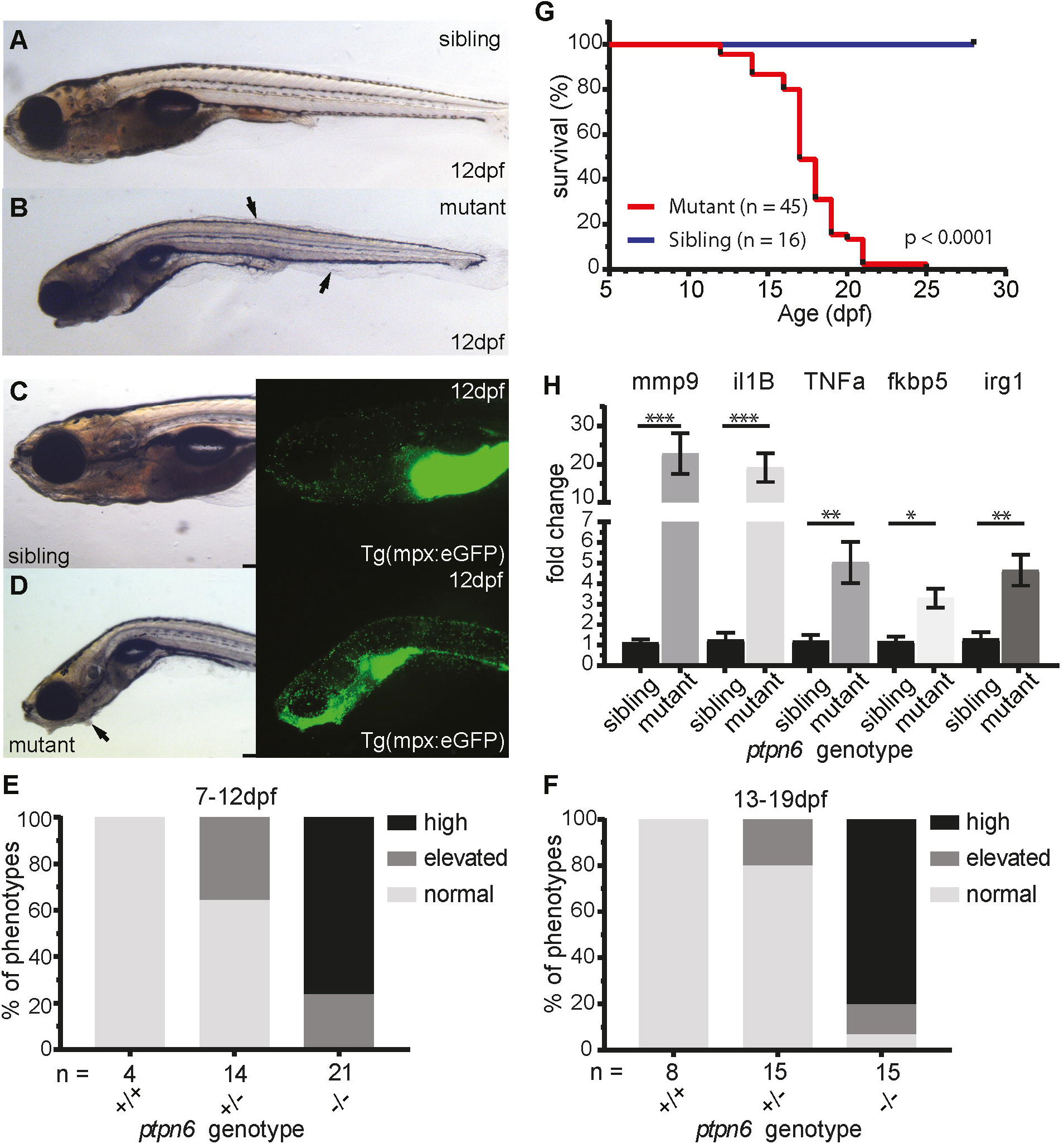
Zebrafish *ptpn6* mutants showan inflammatory response with neutrophil accumulation and do not survive to adulthood. (A-D) Stereo microscope representative pictures of embryos at 12 dpf. (A,C) siblings and (B,D) *ptpn6*^*-/-*^ mutant displaying the “motheaten” like phenotype. Arrows indicate the affected epithelium. (C,D) GFP-positive neutrophils in *tg(mpx:eGFP)* transgenic line localize to the gill and mandibular area in *ptpn6*^*-/-*^ embryos. (E,F) Scoring of phenotypes in age cohorts of larvae. Embryos were classified as normal (distribution and number of neutrophils as in WT larvae), elevated (mildly increased number of neutrophils in gill and mandibular area) or high (massive increase of neutrophil numbers in the gill and mandibular area). (G) Survival curve of *ptpn6* mutants. Curve comparison by log-rank (Mantel-Cox) test. (H) Increased expression of pro-inflammatory genes in mutants. RNA was extracted from larvae sacrificed upon observation of a severe phenotype and their age-matched siblings. Gene expression was determined by qPCR and fold changes were calculated per time point (10, 14 & 18dpf) and pooled afterwards. Statistical comparisons were performed by non parametric ANOVA (Kruskal-Wallis) followed by multiple comparisons (original FDR method Benjamini Hochberg). *p<0.05, **p<0.01, ***p<0.001, error bars = SEM

In accordance with the *Ptpn6*^*me/me*^ and *Ptpn6*^*me-v/me-v*^ mice, *ptpn6* mutant zebrafish do not survive to adulthood (Green and Shultz, 1975; Shultz *et al*., 1984). The absence of Shp1 is lethal at late larval stages with a median survival of 17 days (Fig 2G). To investigate whether affected larvae present with inflammation, we collected mRNA from mutants (n=12) and age matched siblings (n=9) at the end points of the mutant larvae at 10, 14 and 18 dpf. We performed qPCR for several pro-inflammatory genes and genes which are upregulated during inflammation (*mmp9, il1B, tnfα, fkbp5* and *irg1/acod1*), of which the first two were previously found to be upregulated in the *ptpn6* morpholino studies in zebrafish embryos (Kanwal *et al*., 2013). All five genes were significantly upregulated compared to siblings (Fig. 2H), indicating that loss of functional Shp1 evokes an inflammatory response.

### The development of erythroid and megakaryocyte lineages are unaffected in ptpn6 mutants

To assess erythropoiesis we performed an O-dianisidine staining and *gata1 in situ* hybridization on 5dpf embryos. O-dianisidine detects heme in hemoglobin and *gata1* marks early erythrocytes (Iuchi and Yamamoto, 1983; Ransom *et al*., 1996). We found no difference in erythrocyte numbers or distribution between *ptpn6* mutants and siblings (Fig. S2A-E). To investigate the megakaryocyte lineage, we quantified thrombocytes, the nucleated equivalents of mammalian platelets in *Tg(cd41:GFP)* embryos at 5dpf by confocal microscopy. In these transgenic embryos, thrombocytes express a high level of GFP and HSPCs a low level of GFP (Kissa *et al*., 2008). After fixation only GFP^high^ cells are visible (Fig. S2F,G). Counting of GFP^high^ cells revealed no difference in thrombocyte numbers between siblings and *ptpn6* mutants (Fig. S2H). Hence, the erythroid and megakaryocyte lineages in zebrafish embryos lacking functional Shp1 appear to be unaffected at 5 dpf.

### Development of the myeloid cell lineage but not early lymphocyte development is affected in ptpn6 mutants

To investigate the development of leukocytes in *ptpn6* mutants we first determined *l- plastin* expression, a pan-leukocyte marker, by whole mount *in situ* hybridization. *L-plastin* expression was higher in the CHT of *ptpn6* mutants, indicating an increase in total leukocyte numbers (Fig 3A,B). To determine which cell type was responsible for this increase we performed *in situ* hybridizations and stainings, and used transgenic lines to quantify cell numbers. First, we investigated the expression of *ikaros* and *rag1* by *in situ* hybridization, to determine whether the development of the lymphoid lineage was affected. *Ikaros* is an early lymphocyte marker and *rag1* is essential for V(D)J recombination in maturing B and T cells (Willett *et al*., 1997, 2001). As expected, *ikaros* expression was detectable in the CHT and the thymus of 5dpf embryos and no differences in expression level or location were observed between *ptpn6* mutants and siblings (Fig 3C-F). Quantification of the *rag1-positive* area in mutants, heterozygous embryos and wild type siblings revealed no difference in thymus size, a direct read-out for developing B- and T-cell numbers in the thymus (Fig 3G-I). We conclude that early development of the lymphocyte system is not affected in *ptpn6* mutants.

**Figure 3.**
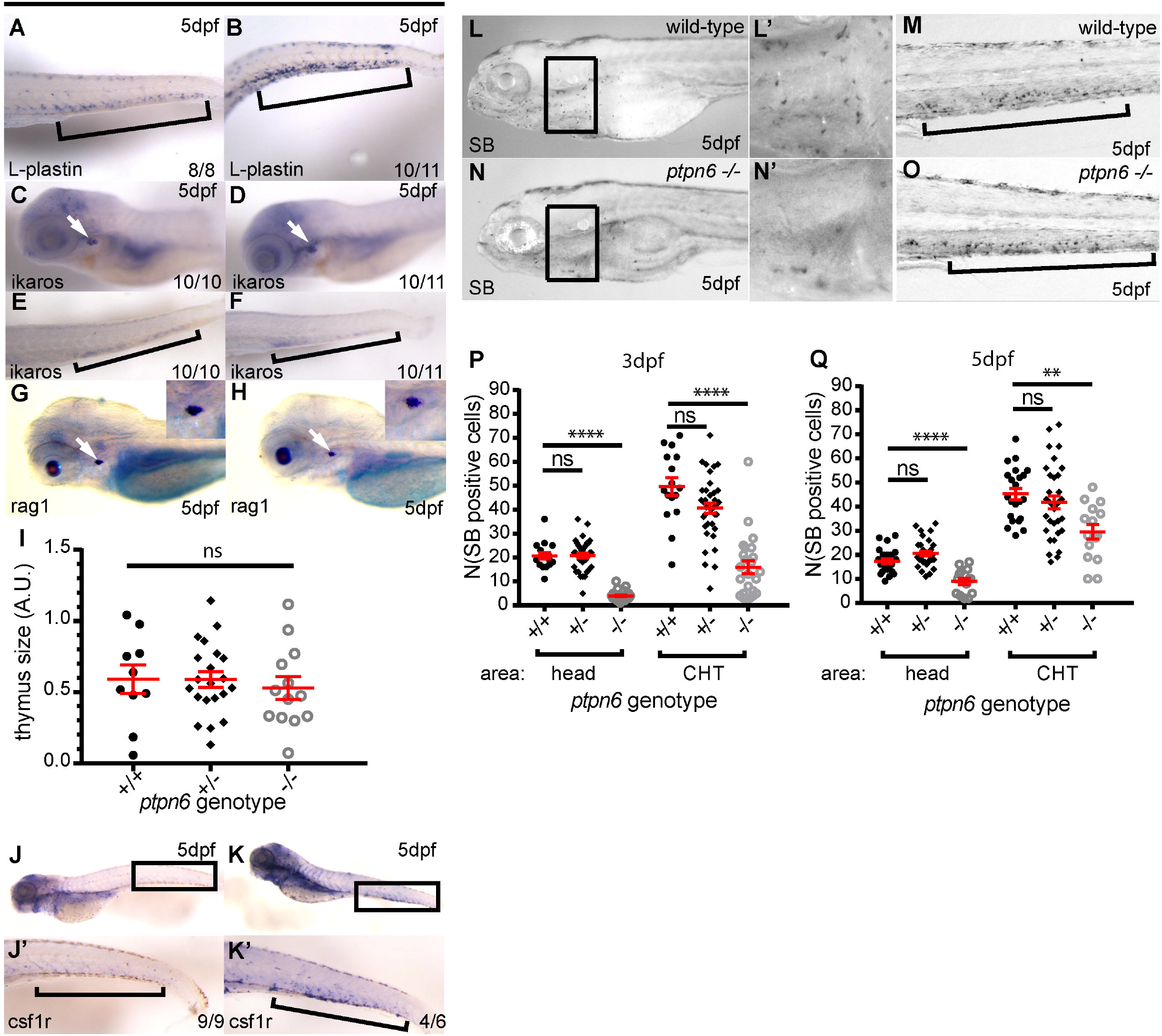
Development of the leukocyte lineages in *ptpn6* mutants. (A-J) A panel of *in situ* hybridization markers for leukocyte lineages was used on 5dpf wild-type and *ptpn6*^*-/-*^ embryos. CHT is indicated by bracket and thymus by white arrow. The number of embryos showing the depicted pattern/ total number of embryos is shown in the bottom right corner. (A,B) L-plastin, pan-leukocyte marker; (C-F) ikaros, lymphoid progenitors, and (G,H) rag1, lymphocytes. Representative lateral views of the thymus are shown, with close ups of the thymus in the inset. (I) quantification of rag1-positive area (n.s., ANOVA). (J,K) csf1r, macrophage marker with (J’,K’) close ups of the CHT. The number of neutrophils in two parts of the embryo was quantified by SB staining at 3 and 5 dpf (L-O). (L’, N’) Show magnifications of the boxed areas in L and N. (P, Q) SB-positive cells were counted in the head (mainly primitive wave) and CHT (mainly definitive wave) at 3 and 5 dpf. Statistical comparisons were performed by ANOVA followed by multiple comparisons (Tukey) for both areas per time point separately. **p<0.01, ****p<0.0001, error bars = SEM

Next, we investigated the two main cell types in the myeloid lineage, the monocytes/macrophages and the neutrophils. Expression of *csf1r*, a marker for macrophages, was upregulated in the CHT of *ptpn6* mutants at 5 dpf (Fig 3J,K). Sudan Black (SB) stains the granules of immature and mature neutrophils and was used to quantify these in 3 and 5dpf embryos. SB-positive cells in the head are mainly derived from the primitive wave at early time points in development, whereas SB-positive cells in the CHT are derived from the definitive wave (le Guyader *et al*., 2008). SB-positive cells were counted in these two areas of the embryos, indicated by a black box (the head, including the heart), and by brackets (the CHT)(Fig 3L-O). Surprisingly, given the late larval phenotype with increased neutrophil numbers (Fig. 2C,D), at 3dpf neutrophil numbers in the head area of *ptpn6* mutants were reduced by approximately 80% compared to WT siblings, and by 65% in the CHT (Fig 3P). At 5dpf these numbers were reduced to approximately 50% and 35% respectively (Fig 3Q). In addition to the SB staining we quantified neutrophils in 3dpf *Tg(mpx:GFP)* embryos, which facilitated counting of neutrophils in intact embryos. Mpx is a neutrophil marker which is expressed already early in neutrophil differentiation, from the promyelocyte phase onwards (Bainton, Ullyot and Farquhar, 1971; Borregaard and Cowland, 1997; Kumar *et al*., 2010). Total *mpx*-positive neutrophil numbers were reduced by 41% in *ptpn6* mutants, compared to wild type siblings. Because *mpx* is expressed from early neutrophil precursors onwards, this indicated that the whole neutrophil lineage was affected, and not just later steps in neutrophil maturation (Fig. S3A,B).

To further investigate the development of the granulocyte-monocyte lineage we used *Tg(mpx:GFP/mpeg1:mCherry)* transgenic fish to simultaneously quantify neutrophil and macrophage numbers. *Mpeg* expression is restricted to the monocyte/macrophage lineage during early development (Ellett *et al*., 2011). By confocal imaging we acquired images of the rostral part and the CHT of 3 and 5dpf *ptpn6* mutant and sibling embryos (Fig 4A-D, 3dpf not shown). Macrophage (Fig 4E) and neutrophil (Fig 4F) numbers in both areas and at both time points were determined using ImageJ. Macrophage numbers were increased in both areas and at both time points during development, whereas neutrophil numbers were reduced. The increase in *mpeg* signal was very pronounced in the area of the pro-nephros, the site of late-larval and adult hematopoiesis in zebrafish (Al-Adhami and Kunz, 1977; Willett *et al*., 1999), where macrophage numbers were up by ∼55%. Similarly, whereas peripheral neutrophils were scarcely found in the head, the pro-nephros was well populated with *mpx*-positive cells.

**Figure 4.**
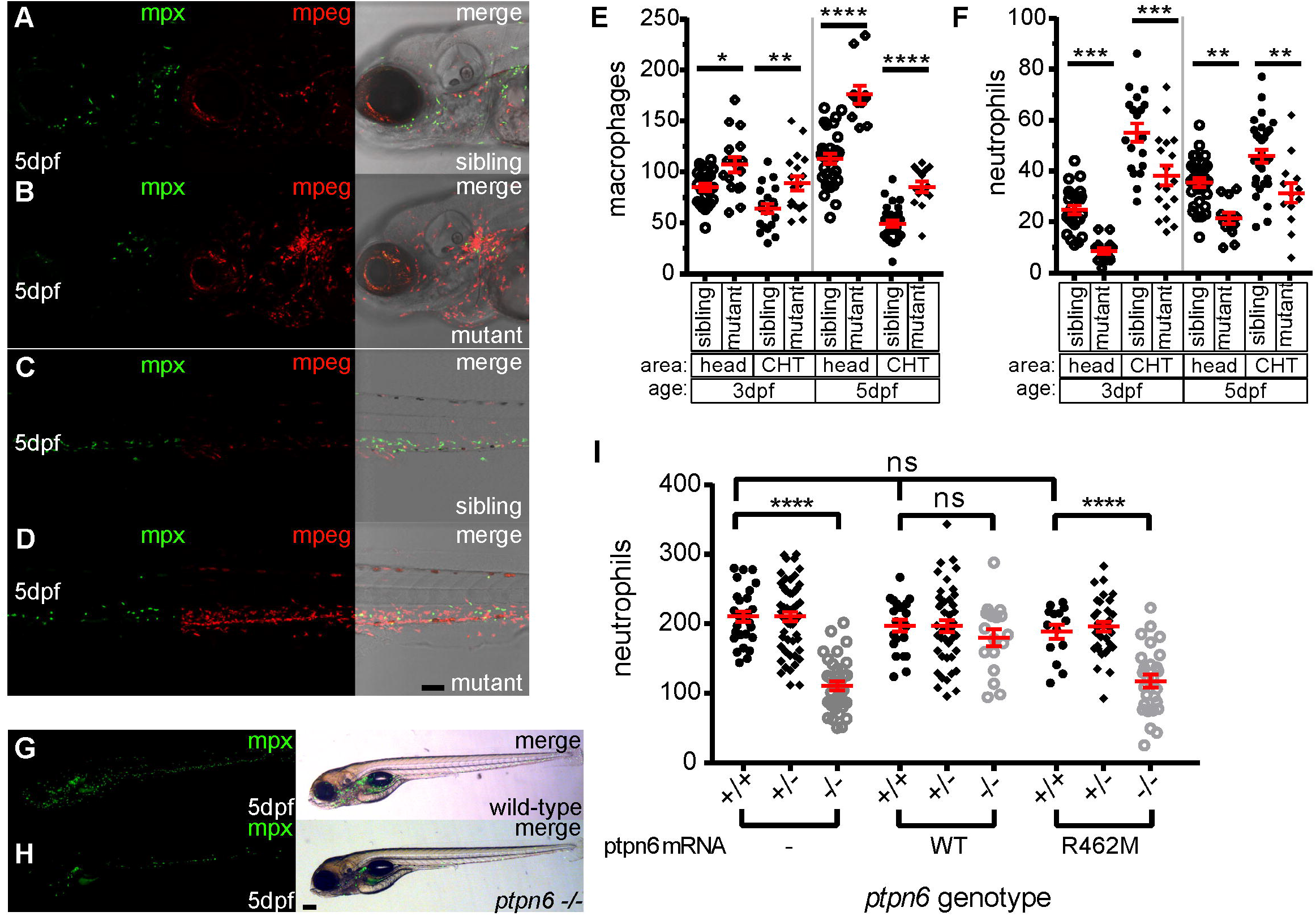
Zebrafish *ptpn6* mutants have an increased number of macrophages and a reduced number of neutrophils, and this phenotype is dependent on phosphatase activity of Shp1. (A,B) Representative pictures of the head area and (C,D) of the anterior part of the CHT of 5dpf *tg(mpx:GFP/mpeg:mCherry) ptpn6* mutant and sibling embryos showing neutrophils and macrophages. 20x objective, pinhole 2AU, step size 1,83μM. (E,F) Scatterplot of macrophage (E) and neutrophil (F) numbers per embryo in the head and CHT of 3 and 5 dpf embryos. Quantification was performed in ImageJ by particle analysis. Statistical comparisons by ANOVA and Sidak’s multiple comparison test for preselected columns. (G,H) Stereo images of 5dpf *tg(mpx:GFP)* wild-type and *ptpn6*^*-/-*^ non injected embryos. Images of WT Shp1 and Shp1- R462M injected embryos are shown in Fig S4. (I) Quantification of total number of neutrophils in *tg(mpx:eGFP)* embryos injected with mRNA encoding WT Shp1 or Shp1-R462M. Scatterplot of total neutrophil numbers at 5dpi per embryo. Quantification was performed in ImageJ by particle analysis. Statistical comparisons by ANOVA and Tukey multiple comparisons test. Scale bar represents (A-D) 100μm and (G-H) 200μm. *p<0.05, **p<0.01, ***p<0.001, ****p<0.0001, error bars = SEM

To exclude apoptosis as the cause of the reduction in neutrophils in *ptpn6* mutant embryos, we performed acridine orange staining. No increase in apoptosis was observed in the CHT of 5dpf *ptpn6* mutant embryos (Fig. S4). These results suggest that *ptpn6* mutant fish suffer from an altered lineage balance within the myeloid cell lineage and early onset increased proliferation in the macrophage lineage.

### The early developmental phenotype of the myeloid lineage is dependent on catalytic activity of Shp1

Next, we questioned whether restoring Shp1 expression rescued the phenotype observed in *ptpn6* mutant embryos. We micro-injected *Tg(mpx:GFP)* embryos at the 1-cell stage with wild type *ptpn6* mRNA and allowed them to develop until 5dpf. The mRNA also encoded GFP, linked with a P2a autocatalytic cleavage sequence. Upon expression, GFP was visible until approximately 3dpf, which allowed us to successfully select injected embryos at 1dpf. Non-injected controls are depicted in figure 4G and H. Expression of wild type Shp1 restored neutrophil numbers in *ptpn6* mutant embryos to those found in non-injected wild type controls at 5 dpf (Fig 4I, Fig. S5). Although catalytic activity is important for Shp1 function, other domains may exert alternative functions, e.g. scaffolding by SH2 domains (Timms *et al*., 1998; An *et al*., 2008; Abram and Lowell, 2017). To investigate whether catalytic activity of Shp1 is essential for the development of neutrophils we injected the catalytically inactive mutant Shp1- R462M. We used the R462M mutant and not the classic catalytic cysteine mutant C456S to avoid inadvertent substrate-trapping effects (Hale and den Hertog, 2018). Upon micro-injection of mRNA encoding GFP-P2a-Shp1-R462M, no increase in total neutrophil numbers was observed in *ptpn6* mutant embryos (Fig 4I, Fig. S5). Expression of Shp1 or inactive Shp1- R462M did not affect the number of neutrophils in wild type or heterozygous siblings (Fig. 4I). Therefore, we conclude that catalytic activity of the PTP domain of Shp1 is essential for its function in the development of the myeloid lineage.

### Early hematopoiesis is affected in ptpn6 mutants

Earlier reports indicated that hematopoietic stem cells (HSCs) in *motheaten* mice are not affected by SHP1 deficiency (Shultz, Bailey and Coman, 1987). However, recently, SHP1 was found to be involved in HSC quiescence in *Scl-CreER/Shp1*^*fl/fl*^ mice (Jiang *et al*., 2018). Therefore we investigated whether HSPC emergence, homing and proliferation was affected in *ptpn6* mutant zebrafish. First, the emergence and early homing of HSPCs at ∼36hpf was investigated by *in situ* hybridization of *c-myb* mRNA in *ptpn6* mutants and siblings (Fig 5A,B). A minor decrease in signal was observed in *ptpn6* mutants. Subsequently *Tg(cd41:eGFP/kdrl:mCherry)* fish were used to quantify HSPCs in the CHT after arrival (Kissa *et al*., 2008; Choorapoikayil *et al*., 2014). Kdrl-positive embryos were selected at 46hpf, and subjected to immunohistochemistry with a GFP specific antibody. Confocal imaging was used to capture the complete CHT, based on the kdrl-mCherry signal (Fig 5C,D). GFP-positive cells in the CHT were quantified and at 46 hpf, a significant reduction in HSPCs in the CHT was observed in *ptpn6* mutant embryos, compared to heterozygous embryos (P<0.05) and a 17% reduction compared to wild type siblings (p<0.01) (Fig 5I). Next, live imaging was performed of the CHT of 5dpf *Tg(cd41:eGFP/kdrl:mCherry) ptpn6* mutant and sibling embryos (Fig 5E,F). The number of GFP^low^ HSPCs was quantified, showing expansion of HSPCs in the CHT over time from 46hpf to 5dpf. The number of HSPCs was 58% higher in mutant embryos at 5 dpf than in heterozygous and wild type embryos (p<0.0001) (Fig 5J). To investigate the cause of the increase in HSPC number, proliferation of HSPCs was assessed in the CHT of *ptpn6* mutant embryos by immunostaining using pHis3- and GFP-specific antibodies of 5dpf *Tg(cd41:eGFP/kdrl:mCherry)* embryos (Fig 5G,H). The *kdrl*-mCherry signal was used to determine which cells were located in the CHT and the pHis3-positive cells were counted. A 34% increase in proliferating cells was detected in the CHT of *ptpn6* mutant embryos compared to sibling cell numbers (p<0.01) (Fig 5K). Taken together, these results suggest that early HSPC emergence and/or arrival in the CHT is reduced or delayed in *ptpn6* mutant embryos and that the proliferation of these definitive blood cell progenitors is enhanced once they seed the CHT.

**Figure 5.**
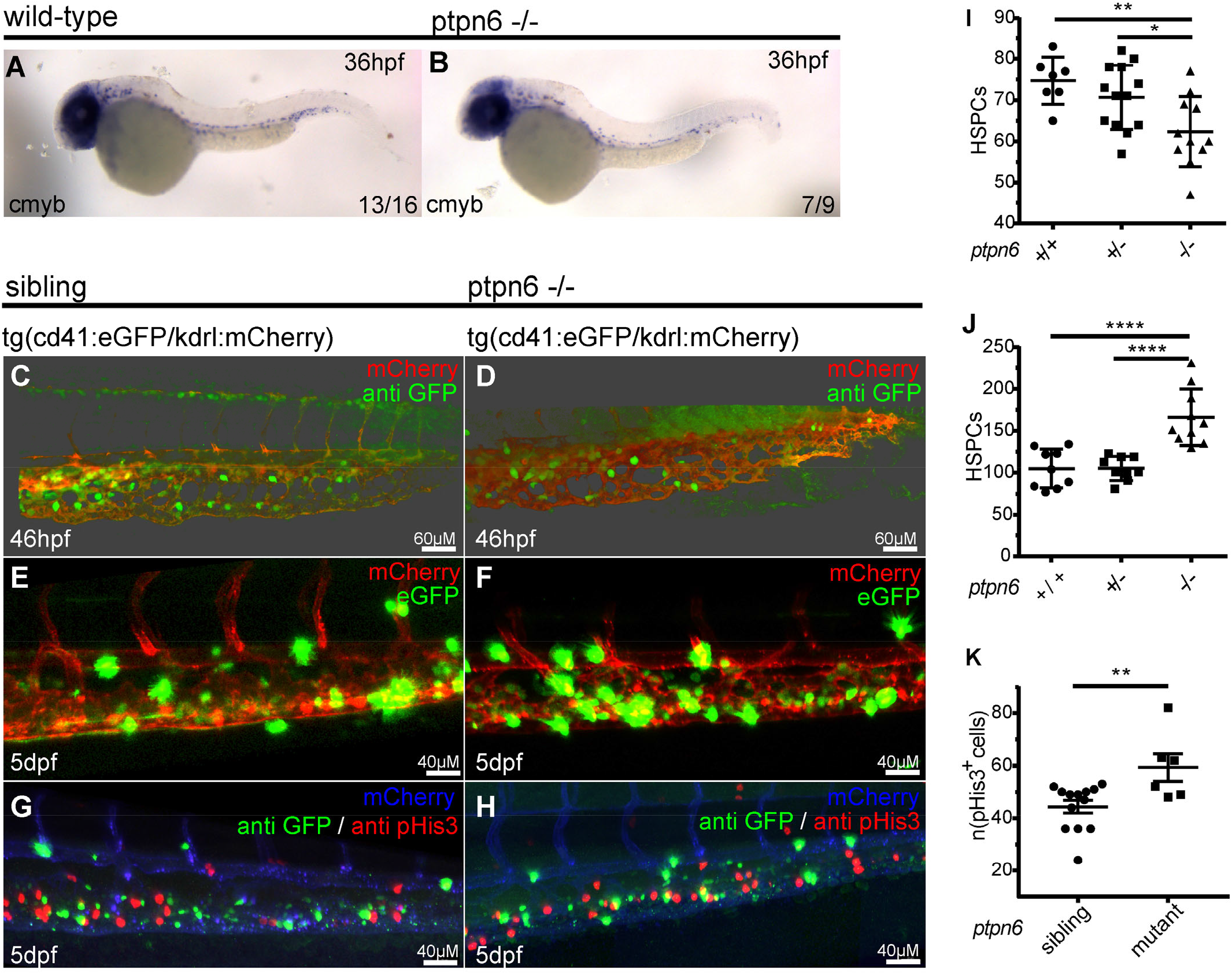
HSPCs in *ptpn6* mutant embryos. (A,B) Stereo images of cmyb *in situ* staining, a HSPC marker, at 36hpf. The number of embryos showing the depicted pattern/ total number of embryos is shown in the bottom right corner. (C-H) confocal images of the CHT of *tg(cd41:eGFP/kdrl:mCherry)* zebrafish, 40x objective, pinhole 2AU. (C,D) Embryos were selected and fixed at ∼46hpf, when no GFP^high^ cells marking thrombocytes were present yet, and subsequently whole-mount immunohistochemistry was performed using a GFP specific antibody. Representative 3D blended rendering images of the CHT of 46hpf embryos. Step size 1.5μm (E,F) Live imaging of 5dpf embryos. Representative 3D maximum intensity projection rendered images of part of the imaged region of the CHT are shown. Step size 2μm. (G,H) 5dpf embryos were fixed and whole-mount immunohistochemistry was performed using a pHis3 and GFP specific antibody. Representative 3D maximum intensity projection images of part of the imaged CHT are depicted. Step size 2μm. (I-J) Scatterplots of quantification of cells in confocal images using IMARIS. (I) GFP positive HSPCs in the CHT of 46hpf embryos and (J) GFP^low^ HSPCs in the CHT of 5dpf embryos. Statistical comparisons were performed by ANOVA followed by a Tukey’s multiple comparisons test. (K) quantification of pHis3-positive cells in the CHT of 5dpf embryos. Means were compared using a 2-sided student’s t-test. *p<0.05, **p<0.01, ***p<0.001, ****p<0.0001, error bars = SEM

### Directional migration of neutrophils and macrophages in response to tailwounding is affected in ptpn6 mutants

Neutrophils and macrophages play an essential role during injury response. They are among the first cells to be recruited to an injury site, where they remove debris and release molecules that promote inflammation and vasodilation. We investigated whether the behavior of neutrophils and macrophages in *ptpn6* mutant embryos differed from siblings by live-imaging of 4dpf embryo tails following amputation of the tip of the tail fin fold (Fig 6A; representative movies are presented in Supplemental Material, SM1 and SM2). The number of neutrophils that migrated into the wound area (closer than 200 μm to the cut site) in mutant embryos was not significantly different from their siblings (Fig 6B). However, the meandering index (net distance travelled/ total distance travelled) of neutrophils outside the wound area was lower for mutant embryos than their siblings (p<0.0001). Speed and wound persistence of neutrophils were not different in mutants lacking functional Shp1 (Fig. 6C).

**Figure 6.**
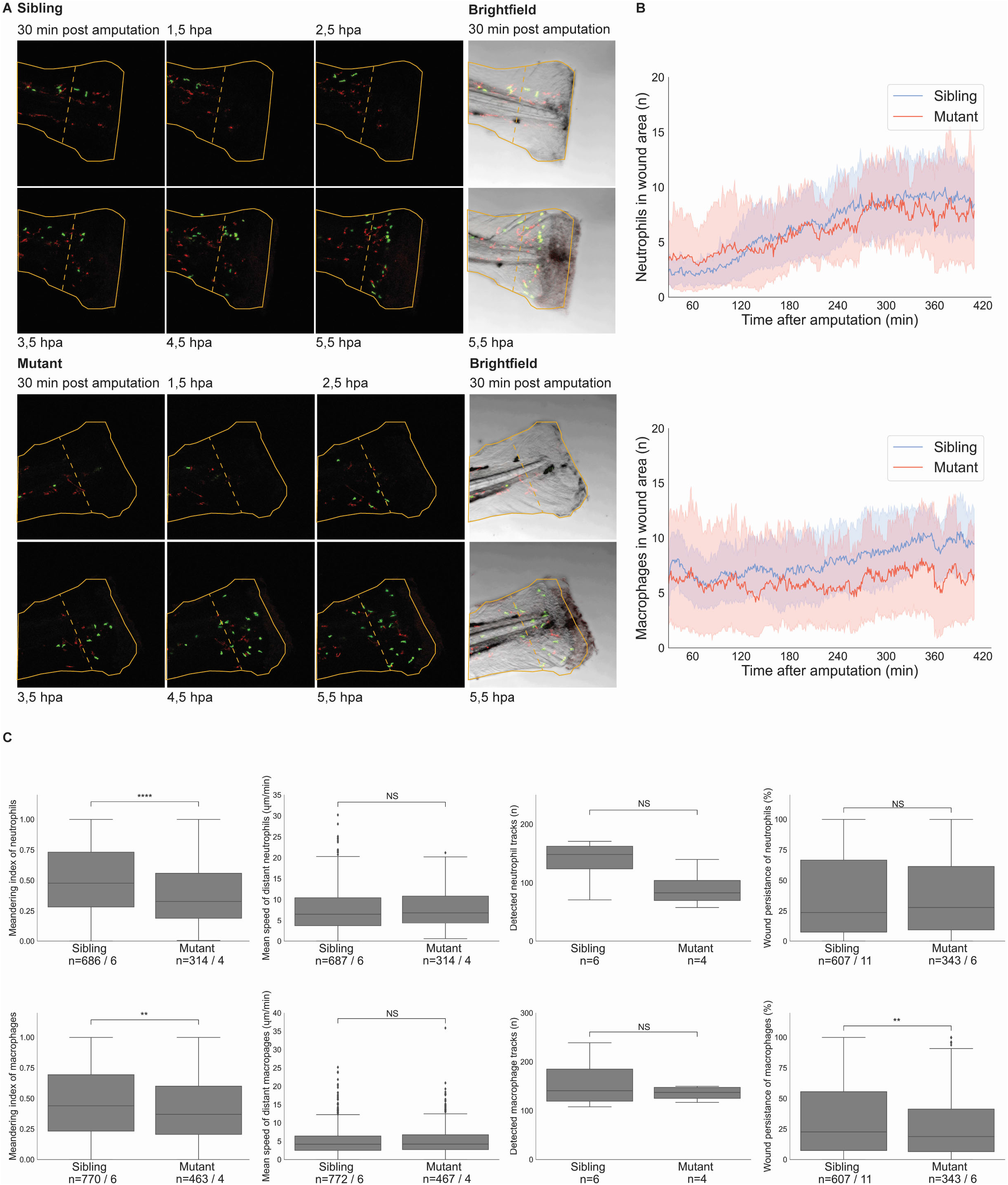
Neutrophil and macrophage migration is affected upon tail fin fold amputation in Shp1 mutants. (A) Stills from live imaging of mutant and sibling 4dpf embryos in *tg(mpx:GFP/mpeg:mCherry)* background show neutrophils in green and macrophages in red. Embryos were imaged every minute for 7 h, see Supplementary Material, SM1 and SM2. The outline of the amputated at the start of imaging is indicated with a solid orange line. The border of the wound area is indicated with a dashed orange line. (B) Cells were detected using Trackmate. Cells within 200 μm of the wound were counted in every frame from 30 min after amputation till 410 min after amputation. Shaded area represents results of bootstrapping. Results of 6 mutants and 11 siblings were analyzed and compared by ordinary least squares regression (statmodels). For neutrophils, p-value=0.78 & 95% CI coefficient = [-2.10 - -1.54]; for macrophages, p-value<0.0001 & 95% CI coefficient = [-0.24 - 0.02]. (C) Meandering index and mean speed were determined for the distant cells (further than 200 μm from the wound edge) of 6 siblings and 4 mutant embryos. Meandering index was defined as the net distance travelled/ total distance travelled of a track. All results of the measurements in (C) were compared using unequal variance T-test (SciPy) and two sided 95% confidence intervals (statmodels). Meandering index was significantly reduced for neutrophils, p-value<0.0001, 95% CI[0.48- 0.52] vs [0.36-0.42] and macrophages, p-value=0.0014, 95% CI[0.39-0.44] vs [0.45-0.49]. Mean speed was not significantly different for neutrophils, p-value=0.5, 95% CI[7.3- 8.1] vs [7.4-8.5] and macrophages, p-value=0.48, 95% CI[4.82-5.35] vs [4.89-5.61]. The number of tracks was not significantly different for the entire tail of 6 siblings and 4 mutant embryos: neutrophils, p- value=0.092; macrophages, p-value=0.37. Wound persistence was determined for all cells that entered the wound area of 11 siblings and 6 mutants. For neutrophils, p-value=0.78, 95% CI[0.35- 0.41] vs [0.35-0.42]; macrophages, p-value=0.014, 95% CI[0.33- 0.38] vs [0.28-0.34]. NS, not significant; * p<0.05, ** p<0.01, *** p<0.001, **** p<0.0001. Boxplots show quartiles of data. Outliers were determined as datapoints outside 1.5 interquartile range. Number of cells and number of embryos are mentioned as n= cells/ embryos.

The total number of macrophages in embryos lacking Shp1 is increased at 3 dpf and 5 dpf (Fig. 4E). In contrast, the number of macrophages in the wound area is significantly decreased in mutant embryos compared to their siblings (p<0.0001) (Fig 6A,B, Supplemental Material SM1 and SM2). This is supported by the notion that wound persistence of macrophages is decreased in mutants compared to siblings (p=0.007) (Fig 6C). The meandering index of macrophages distant from the wound was significantly reduced in mutant embryos (p=0.001), while the speed was not affected (Fig 6C). In contrast to the total number of neutrophils and macrophages (Fig. 4E,F), the number of macrophages and neutrophils that were detected in the tail was not significantly different between mutants and siblings, although the number of neutrophils in mutant embryo tails trends towards a reduction (Fig 6C). Despite the lower number of macrophages in the wound area, there was no difference in regeneration between *ptpn6* mutant embryos and their siblings (Fig S6). Taken together, embryos lacking functional Shp1 exhibited differences in behavior of neutrophils and macrophages, which however did not affect caudal fin fold regeneration.

## DISCUSSION

SHP1 *null* mutations have long been studied in the context of auto-immunity and inflammation in the *motheaten* mouse. Because mice develop in utero, and *motheaten* mice are already severely affected at birth (Green and Shultz, 1975) it has been difficult to study the early development of the hematopoietic system and inflammatory phenotype. In this study we developed a zebrafish model to study hematopoiesis during embryonic development in the absence of Shp1. In the mouse embryo, HSCs emerge from the dorsal aorta and colonize the liver before populating the adult organs of hematopoiesis (Sánchez *et al*., 1996; de Bruijn *et al*., 2000; Kumaravelu *et al*., 2002; Boisset *et al*., 2010). An equivalent process takes place in zebrafish embryos, where HSPCs emerge from the ventral wall of the dorsal aorta, temporarily colonize the CHT, which is homologous to the mouse fetal liver, before migrating to the head kidney, the adult site of hematopoiesis in zebrafish (Kissa and Herbomel, 2010; Murayama *et al*., 2006). Furthermore, the zebrafish adaptive immune system is not functionally mature until 4-6 weeks post fertilization, allowing analysis of the function of Shp1 in innate immunity during embryonic development.

Shp1 knock-out in zebrafish is lethal at late larval stages, but did not induce obvious morphological defects in embryos. Morpholino-mediated knockdown of Shp1 induced pleiotropic defects from 3 dpf onwards (Kanwal et al., 2013). The difference between the morpholino-mediated knockdown and the knockouts we generated may be caused by intrinsic differences between morpholino-mediated knockdown and genetic knockouts. Morpholinos block translation or splicing from the moment the morpholino is administered, i.e. the one-cell stage, whereas in the genetic knockdown, maternally contributed Shp1 may persist, which may result in a later onset of the phenotype. It is interesting to note that the Shp1 morpholino elicited skin lesions similar to the lesions observed in the genetic mutant (Fig. 2A-D). The inflammatory response was also common between the morpholino-mediated knockdown and the genetic knock-out, suggesting that the response to loss of functional Shp1 was similar, but that the timing was different.

Mutant zebrafish larvae present with hyperinflammation, characterized by strong upregulation of several inflammatory genes. The phenotype includes skin lesions and infiltration of gills and mandibular area with neutrophils. The hyperinflammation, the skin lesions and the gill infiltration by neutrophils are very similar to the *motheaten* appearance and lethal pneumonitis found in *motheaten* mice (Green and Shultz, 1975; Jiao *et al*., 1997). It also supports the sporadic cases of humans that present with mutations in *ptpn6*, which have been associated with neutrophilic dermatoses, and emphysema (Nesterovitch *et al*., 2011; Bossé *et al*., 2019). The neutrophil specific Shp1 knock-out mouse line suggests that the skin lesions and inflammation are caused by mutant neutrophils. However, the lethal pneumonitis is not recapitulated by neutrophil-specific mouse knock-out lines, which indicates that other cell types are involved in causing this phenotype. The strong occurrence of gill and mandibular infiltration in Shp1 mutant zebrafish composes an ideal model to investigate the development of lethal pneumonitis/gill infiltration due to Shp1 knock-out.

Shp1 deficiency led to multiple hematopoietic defects during embryonic development, affecting the HSPCs and myeloid cell lineages. The number of emerging HSPCs was reduced in mutants compared to siblings. However, this reduction was compensated by enhanced proliferation of HSPCs later during development. These results are in contrast with a previous report that showed no effect on HSPCs in *motheaten* mice (Shultz, Bailey and Coman, 1983). However, our results support the finding that knock-out of Shp1 in HSPCs causes enhanced proliferation in mouse embryos (Jiang *et al*., 2018). A possible explanation for this apparent discrepancy is that the earlier report missed an effect on HSPCs due to the use of a spleen assay to determine the number of HSPCs. Enhanced proliferation of HSPCs may have compensated for the lower number of early HSPCs. The zebrafish model facilitated continuous analysis of hematopoiesis from the moment HSPCs emerged from the dorsal aorta, thus revealing reduced numbers of HSPCs at the start, which showed enhanced proliferation later during development (Fig. 5).

The number of macrophages was increased in *ptpn6* mutant zebrafish embryos (Fig. 4), which is consistent with earlier findings in *motheaten* and *motheaten viable* mice. Macrophages from the spleen of *motheaten* mice showed increased proliferation and faster maturation than wild-type macrophages *in vitro* (McCoy *et al*., 1982, 1983). Tissue sections from *motheaten viable* mice also show a significant increase in macrophage numbers in spleen, bone marrow and peripheral tissues (Nakayama *et al*., 1997). Finally, ES cells expressing dominant negative Shp1 differentiate into a strongly increased number of myeloid cell colonies in a hematopoietic colony forming assay (Paling and Welham, 2005). However, the observation that the increase in macrophage number occurred at the expense of the number of neutrophils (Fig. 4) has not been noticed in mice. It would be interesting to investigate the early numbers of neutrophils in mice lacking functional Shp1 to verify if the phenomenon we noticed in zebrafish is similar in mice.

Directional migration of neutrophils and macrophages after tail fin fold wounding was reduced (Fig. 6, Supplemental Material SM1 and SM2). Furthermore, the number of macrophages that were attracted to the wound site was reduced. This shows that Shp1 has a role in the attraction-migration process that occurs directly after wounding. It is likely that Shp1 has a role in sensing the attraction signal, rather than in sending the signal, given the predominant expression of Shp1 in hematopoietic cells and given the role of Shp1 in intracellular signalling (Plutzky, Neel and Rosenberg, 1992; Abram and Lowell, 2017). It is interesting to note that, even though the meandering index of neutrophils is reduced, this did not lead to a reduction in the number of neutrophils that reached the wound site. Apparently, there is still a strong attraction of neutrophils to the wound site. However, the cells struggled to migrate efficiently towards their target. The lower number of macrophages that were present at the wound site indicated that less macrophages were attracted to the wound site, which was surprising because the number of macrophages in embryos lacking functional Shp1 was enhanced (Fig. 4). However, no increase in macrophage number was seen in the tails imaged for the tailwounding experiments (Fig. 6). This indicates that the macrophages were primarily located in the anterior of the embryos, instead of the tail region. Further research is needed to definitively establish the effect of Shp1 on macrophage migration.

Furthermore, regeneration was unaffected in *ptpn6* mutant zebrafish embryos (Fig. S6), despite the lower number of macrophages that reached the wound site. This is surprising, given a previous report that showed that ablation of macrophages severely impaired regeneration (Li *et al*., 2012). Possibly the low number of macrophages that reached the wound site was still sufficient for normal regeneration. Another possibility is that Shp1 knock-out has affected signalling in macrophages in such a way that fewer macrophages were required for the regeneration response. Taken together, whereas Shp1 is dispensable for regeneration of the caudal fin fold, Shp1 is required for normal behaviour of neutrophils and macrophages. More research is needed to determine the exact role of Shp1 in neutrophil and macrophage function. The *ptpn6* mutant we generated will be instrumental for further research into the function of Shp1 in neutrophils and macrophages.

## MATERIALS AND METHODS

### Zebrafish Husbandry

All fish were housed and handled according to local guidelines and policies in compliance with national and European law. All procedures involving experimental animals were approved by the local animal experiments committee (AVD8010020173786). Larvae raised in survival experiments were fed shrimp larval diet with a diameter of 5-50μM (Caviar, Bernaqua) to control for the possible inability of small larvae to eat the normal feed.

### Generation of the *ptpn6* mutant

The gRNA was designed using the CHOPCHOP web tool for genome editing (https://chopchop.cbu.uib.no/) and produced according to the protocol of (Gagnon *et al*., 2014)) using the published constant reverse oligo and the following gene-specific forward primer: 5’taatacgactcactataggaaccctacaggataaagagttttagagctagaaatagcaag3’. Ribonucleoprotein complexes of n-terminal GFP labeled Cas9 protein and the gRNA were generated by mixing Cas9 protein and 75 ng gRNA in 4 μl buffer. The complex was injected into wild type TL embryos at the one-cell stage. GFP positive embryos were selected at 6hpi and raised to adulthood. F0 was screened for germline mutations by PCR genotyping of the offspring for heterozygous mutations. A 7bp mutation was identified and the founder was outcrossed twice to TL to obtain a stable line. PCR products of the offspring were sub-cloned in pBluescript sk^−^ for sanger sequencing of the mutation.

### Genotyping

*Ptpn6* mutations were analyzed by PCR of lysed embryos or fin clips using the following primers targeted to the mutation site: fw 5’ggattcaaaacacaggggatta3’ and rev 5’tttaacttggcaaacacacctg3’. PCR products were run on a 4% agarose gel for analysis.

### Polyclonal zebrafish Shp1 antibody

GST-Shp1 fusion protein was produced by expression of pGex-GST-Shp1 in *E. coli* B12 followed by glutathione-agarose affinity purification (Sigma). Purified protein was shipped for antibody production (custom polyclonal antibodies, Eurogentec). Rabbit polyclonal anti-Shp1 was purified in house by affinity purification. Briefly, serum was precleared using immobilized GST and GST-fusion protein of highly homologous zebrafish Shp2, to remove antibodies binding GST and antibodies cross reacting with Shp2. Subsequently, Shp1-specific antibodies were affinity purified using immobilized GST-Shp1 fusion protein. Before use, zfShp1-specific antibody (1:500 in 5% milk) was incubated with purified zfShp2 protein for 1 h to prevent residual cross reaction with Shp2.

### Immunoblotting

Prior to snap freezing, fin clips of 5dpf embryos were collected for genotyping. 3 embryos/genotype were pooled and lysed in cold lysis buffer (25mM HEPES pH 7.4, 150mM NaCl, 0.25% deoxycholate, 1% triton X-100, 10mM MgCl_2_, 1mM EDTA, 10% glycerol + 1:10 cOmplete mini protease inhibitor cocktail (Roche)) by incubation on ice followed by sonication (Bioruptor, 15 min, 30 sec on/off). Lysate of HEK2933T cells expressing zebrafish Shp1 was used as a control. Samples were resolved on a 10% acrylamide gel and transferred to a PVDF membrane. Immunoblotting was performed using Akt-specific (1:500, 9272S Cell Signaling) and Shp1-specific (1:500) antibodies.

### Confocal, fluorescence, bright field microscopy

Confocal microscopy was performed on a Leica SP5. Embryos were anesthetized in tricaine and mounted in 0.8% low melting agarose in E3 medium in a glass cover dish. For live imaging mounted embryos were covered in E3 medium containing tricaine. Stereo-fluorescence imaging and bright field imaging of stained embryos and *in situ* hybridizations was performed on a Leica M165FC connected to a DFC420C camera. Images were processed using imageJ or Imaris.

### Quantitative real-time PCR

Larvae were monitored during raising and were sacrificed at a defined end point (curved, skinny, residing at the bottom of the tank). If 2 or more mutant larvae of the same age were available healthy siblings in the same tank were picked and sacrificed to serve as control samples. Sample sizes and larval age of the larvae used in qPCR are listed in Table S1. Total RNA was extracted from genotyped larvae in Trizol (Invitrogen). Snapfrozen samples were crushed with an Eppendorf tube pestle and further homogenized using a syringe. cDNA was synthesized using superscriptIII first strand synthesis kit (Invitrogen). qRT-PCR was performed using SYBR green and a CFX-96 Connect Real-Time system (Bio-Rad). Data was analyzed by applying the 2^−ΔΔCt^ method (Livak and Schmittgen, 2001). Primers are listed in Table S2.

### Sudan Black, O-dianasidine & Acridine Orange staining

3 and 5dpf embryos were fixed and stained using Sudan Black (Sigma-Aldrich) (le Guyader *et al*., 2008) and counted as published (Choorapoikayil *et al*., 2014). O-dianisidine (Sigma-Aldrich) stainings were performed as described (Iuchi and Yamamoto, 1983), but embryos were fixed afterwards (4% PFA) and cleared in 2:1 benzylbenzoate/benzylalcohol. Acridine Orange staining was carried out as described by Choorapoikayil (Choorapoikayil *et al*., 2014).

### *In situ* hybridization

Embryos were fixed in 4% PFA in PBS. *In situ* hybridizations were performed as previously described (Thisse and Thisse, 2008). Published Digoxigenin-UTP-labeled riboprobes were used (Choorapoikayil *et al*., 2014).

### Whole mount immunohistochemistry

Embryos were fixed overnight in 2% PFA. Samples were washed 2x in PBS + 0,1% Tween-20 (PBS-T) followed by 15 min permeabilization in 1- μg/ml proteinase K (Roche) and 3 washes in PBS-T. Samples were blocked for at least 2 h with 10% goat serum + 0.3% triton X-100 in PBS and incubated overnight with chicken anti-GFP (1:500, Aves Labs Inc) or rabbit anti-pHis3 (1:250, Merck Millipore). Embryos were washed 10x 10min in PBS + 0.3% Triton X-100 (PBS-X), followed by 1 h blocking and overnight incubation in either anti-chicken Alexa 488 (1:500, Invitrogen) or Cy5 anti-rabbit (1:500, Jackson Immuno Research). The next day samples were repeatedly washed in PBS-X for 3-4 hrs and imaged.

### Plasmid construction, RNA synthesis and micro – injections

The coding sequence of zebrafish Shp1 was obtained by PCR (fw 5’accctgtttacgtgtcgaga 3’, rev 5’agccttggctcagttttctt3’) from a mix of cDNA of 3 and 5dpf TL embryos and cloned into pCS^2+^eGFP-p2a using infusion (Takarabio). The R462M point mutation in the PTP domain was introduced by Q5 site-directed mutagenesis (NEB). pCS^2+^ constructs were linearized using NotI and transcribed *in vitro* (mMessage machine SP6, ambion). mRNA was injected in 1 cell stage embryos at a concentration of 5 ng/μl.

### Tailwound assays

4 dpf *Tg(mpx:GFP/mpeg:mCherry)* embryos were anesthesized with tricaine and tails were transsected distal to the notochord by scalpel blade. Embryos were immediately mounted for confocal imaging and imaged from ∼30 min post wounding, for 7 h, every minute. Particles were detected and tracks were analyzed using the Trackmate plugin from Fiji (Tinevez *et al*., 2017). The linear motion algorithm was used with a maximum allowed gap of 4 time frames and a maximum allowed radius of 50 μm. Wound area was defined as 200 μm and closer from the edge of the tail in the first timeframe. Mean speed of tracks outside the wound area was determined by dividing the total distance/ the total time of the track. Meandering index of tracks outside the wound area was determined by dividing the net distance (distance between first and last point of track)/ the total distance of the track. Wound persistence was determined for each track that entered the wound area by dividing the number of time frames spent inside the wound area/ number of time frames left till the end of the movie.

### Statistical analysis

Analytical statistics were performed in GraphPad Prism 7 (version 7.04). For the tailwounding descriptive statistics were determined using Python (matplotlib 3.3.2, numpy 1.18.5, scipy 1.50, statmodels 0.11.1).

### Regeneration assay

Zebrafish embryos were amputated at the tail as previously described (Hale *et al*., 2017). Amputations were performed at 2 dpf and regeneration was analyzed at 5 dpf. Regenerated tails were imaged and whole embryos were lysed for genotyping. Regeneration was quantified by measuring the length from the tip of the notochord to the end of the fin-fold in ImageJ.

## Acknowledgements

The authors would like to thank the animal caretakers at the Hubrecht Institute for excellent care of the zebrafish.

## Author contributions

Conceptualization: P.A.B., M.A., J.d.H.; Methodology: P.A.B., M.A., J.d.H.; Validation: P.A.B., M.A.; Formal analysis: P.A.B., M.A.; Investigation: P.A.B., M.A.; Resources: P.A.B., M.A., H.P.S., J.d.H.; Data curation: P.A.B., M.A.; Writing – original draft: P.A.B., M.A., J.d.H.; Writing - review & editing: M.A., H.P.S., J.d.H.; Visualization: P.A.B., M.A.; Supervision: H.P.S., J.d.H.; Project administration: J.d.H.; Funding acquisition: H.P.S., J.d.H.

