## Supplemental Material for "Loss of Shp1 impairs myeloid cell function and causes lethal inflammation in zebrafish larvae"

**Supplemental Movies**

Imaging of mutant and sibling 4dpf embryos in *tg(mpx:GFP/mpeg:mCherry)* background show neutrophils in green and macrophages in red. Embryos were anesthesized with tricaine and tails were transsected distal to the notochord by scalpel blade. Embryos were mounted immediately for confocal imaging and imaged every minute from ~30 min post amputation onwards for 7 h. Representative movies are shown.

**SM1 – sibling**

**SM2 – ptpn6$^{-/-}$**

**Supplemental Figures**

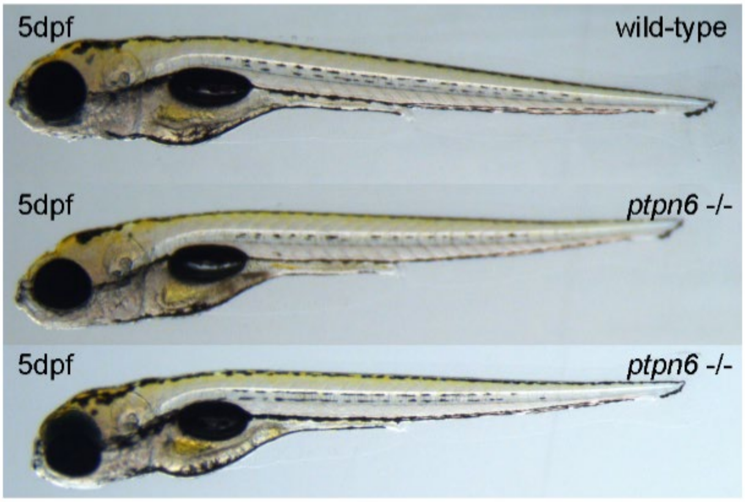


Fig. S1. No morphological defects in 5 dpf embryos lacking functional Shp1. Wild type and *ptpn6*$^{-/-}$ embryos were imaged using a stereomicroscope at 5 dpf. Representative images are shown.

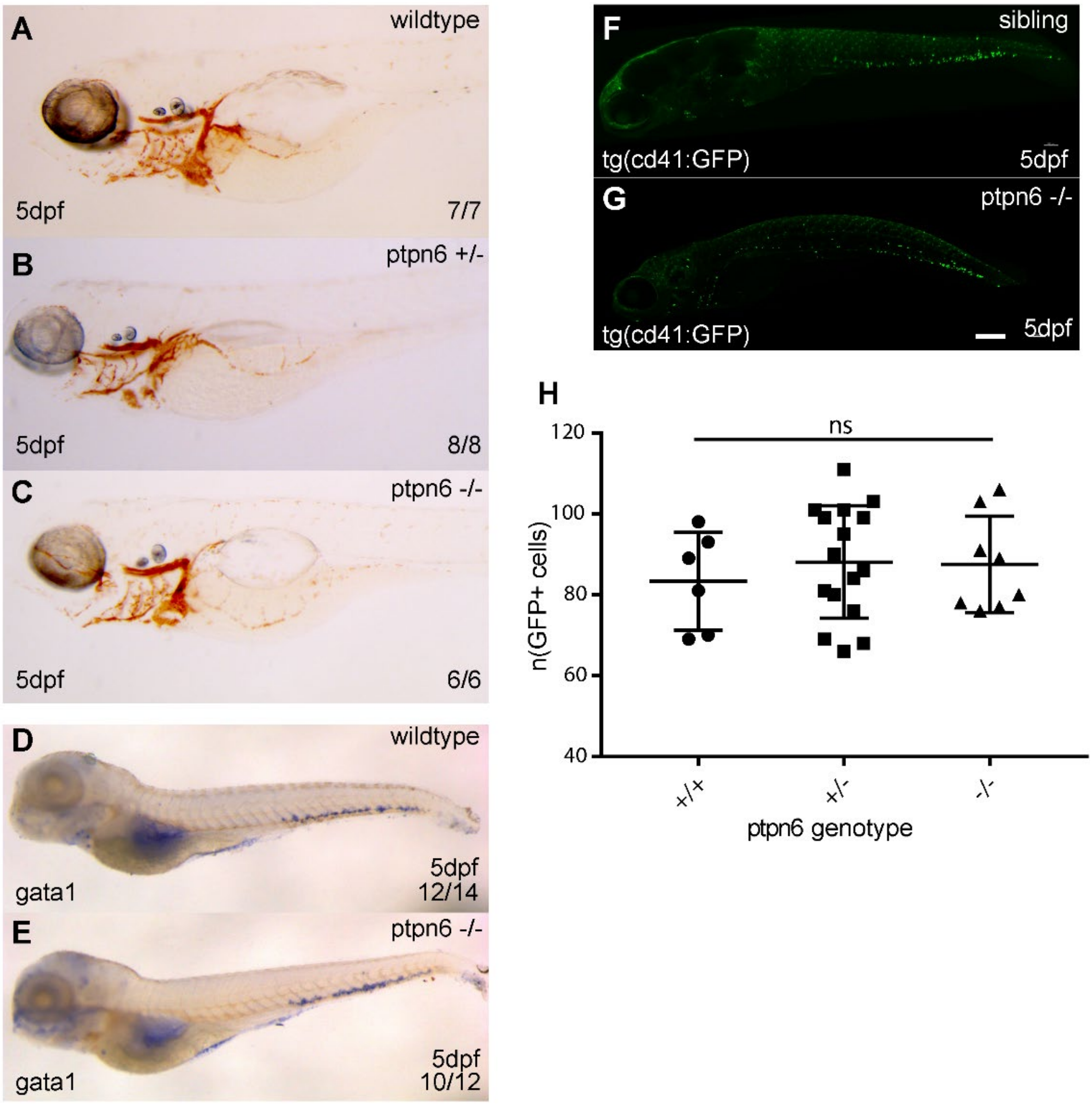


Fig. S2. The erythroid and megakaryocyte lineages are not affected by *ptpn6* knock out. (A-C) Representative stereo images of O-dianasidine stainings of 5dpf embryos and (D,E) *gata1* whole mount in situ hybridization.The number of embryos showing the depicted pattern / total number of embryos is shown in the bottom right corner. (F,G) Representative confocal images of fixed 5dpf *tg(cd41:GFP)* embryos showing thromobyctes, 20x objective. Scale bar represents 200µm. (H) quantification of thrombocytes by IMARIS. Statistical comparisons were performed by ANOVA.

.

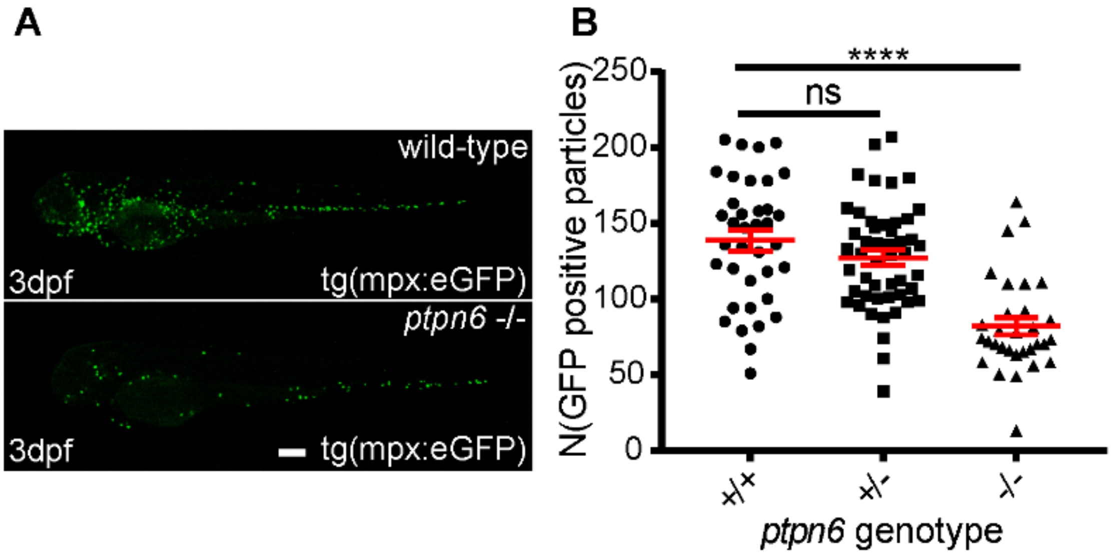


Fig. S3. Reduced numbers of neutrophils in *ptpn6*$^{-/-}$ embryos at 3 dpf. (A) Representative images of wild type and *ptpn6*$^{-/-}$ embryos in *tg(mpx:eGFP)* background, showing reduced number of GFP-positive neutrophils in mutant embryos, lacking functional Shp1. Scale bar represents 200µm. (B) Quantification of the number of GFP-positive neutrophils in wild-type, heterozygous and homozygous *ptpn6* mutant embryos at 3 dpf. Quantification was performed in ImageJ by particle analysis. ANOVA and multiple comparisons (Tukey) were applied for statistical comparisons. ****$p<0.0001$, error bars = SEM

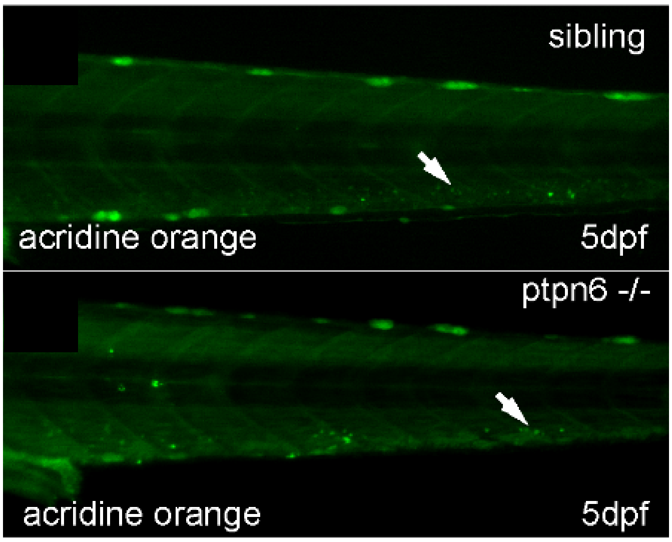


Fig. S4. The reduction of neutrophil number in the CHT of embryos lacking functional Shp1 is not caused by increased apoptosis. Stereo fluorescent images of the CHT of 5dpf embryos stained with acridine orange. Arrows indicate the bright spots representing apoptotic cells. 20x objective, Pinhole 2AU, step size 2.52µm

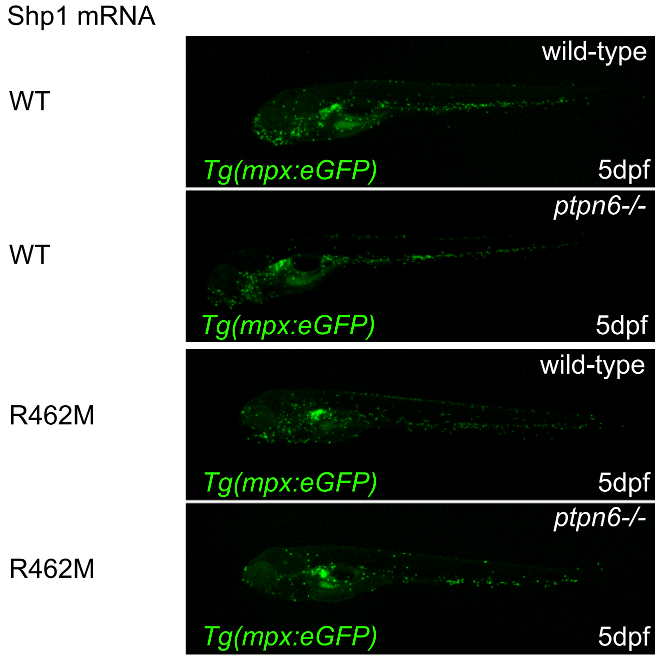


Suppl Fig. 5. Reduced number of neutrophils in *ptpn6*$^{-/-}$ embryos is rescued by expression of Shp1, but not catalytically inactive Shp1-R462M. At 5 dpf, images were obtained of wild type or *ptpn6*$^{-/-}$ embryos in the *Tg(mpx:eGFP)* transgenic background that had been injected at the one-cell stage with synthetic mRNA encoding Shp1 or catalytically inactive Shp1-R462M. GFP-positive neutrophils were quantified and the quantification is depicted in Fig. 4I. Representative images of embryos are depicted here.

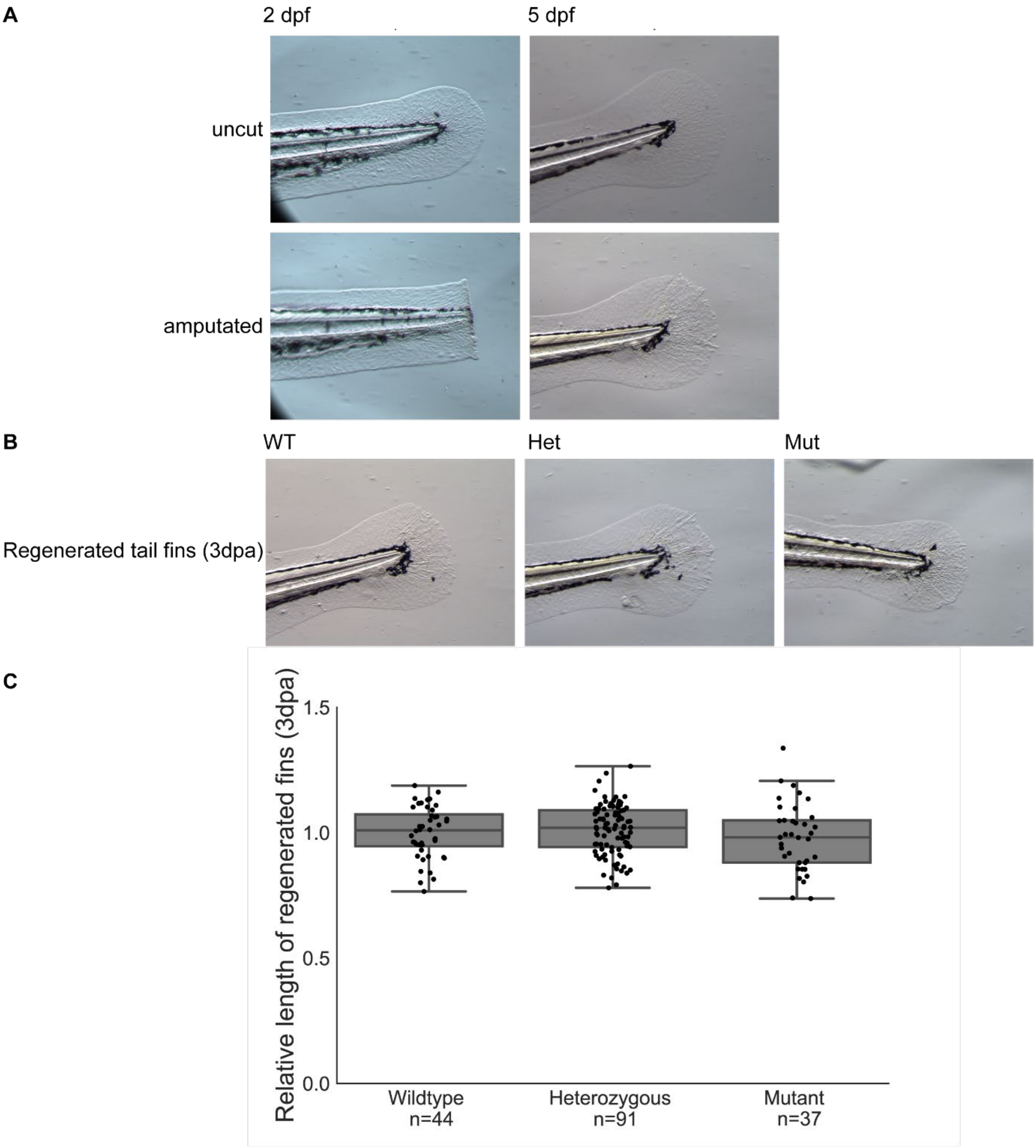


Fig. S6. Tail fin fold regeneration is not affected in Shp1 mutant zebrafish embryos. A. Tail fin folds were cut at the tip of the notochord at 2 dpf. Three days after amputation the tail fin fold regenerated and was indistinguishable from the tail fin fold of uncut controls. Representative pictures are shown. B. There is no difference between WT, heterozygous and mutant Shp1 siblings in tail fin fold regeneration. Representative pictures are shown. C. Length from the tip of the notochord to the end of the fin fold was measured at 3 dpa. Results were normalized to the results of WT siblings. One way Anova resulted in p-value=0.38 (statmodels). Boxplots represent quartiles. All observations are added as single dots.
